# Comparison of TIMS-PASEF quantitative proteomics data-analysis workflows using FragPipe, DIA-NN, and Spectronaut from a user’s perspective

**DOI:** 10.1101/2021.11.29.470373

**Authors:** Alejandro Fernández-Vega, Federica Farabegoli, María M. Alonso-Martínez, Ignacio Ortea

## Abstract

Data-independent acquisition (DIA) methods have gained great popularity in bottom-up quantitative proteomics, as they overcome the irreproducibility and under-sampling limitations of data-dependent acquisition (DDA). diaPASEF, recently developed for the timsTOF Pro mass spectrometers, has brought improvements to DIA, providing additional ion separation (in the ion mobility dimension) and increasing sensitivity. Several studies have benchmarked different workflows for DIA quantitative proteomics, but mostly using instruments from Sciex and Thermo, and therefore, the results are not extrapolable to diaPASEF data. In this work, using a real-life sample set like the one that can be found in any proteomics experiment, we compared the results of analyzing PASEF data with different combinations of library-based and library-free analysis, combining the tools of the FragPipe suite, DIA-NN and including MS1-level LFQ with DDA-PASEF data, and also comparing with the workflows possible in Spectronaut. We verified that library-independent workflows, not so efficient not so long ago, have greatly improved in the recent versions of the software tools, and now perform as well or even better than librarybased ones. We report here information so that the user who is going to conduct a relative quantitative proteomics study using a timsTOF Pro mass spectrometer can make an informed decision on how to acquire (diaPASEF for DIA analysis, or DDA-PASEF for MS1-level LFQ) the samples, and what can be expected depending on the data analysis tool used, among the different alternatives offered by the recently optimized tools for TIMS-PASEF data analysis.

## 1. Introduction

Data-independent acquisition (DIA) methods have gained great popularity in bottom-up quantitative proteomics, as they overcome the irreproducibility and under-sampling limitations of data-dependent acquisition (DDA) due to the semi-stochastic nature of this type of acquisition, showing also a superior quantification accuracy than DDA [1,2]. In DIA, MS/MS spectra are acquired for all precursor ions within isolation windows of defined *m/z* ranges. The major complexity lies in deconvoluting these complex MS/MS spectra using specialized software.

The diaPASEF method, recently developed for the timsTOF Pro mass spectrometers (Bruker) [3], makes use of the TIMS (trapped ion mobility spectrometry) module, which allows ion packets to be trapped, stored, and then resolved according to their different ion mobility (process termed PASEF, parallel accumulation serial fragmentation) [4], being subsequently separated by their *m/z* in the quadrupole before being fractionated (MS/MS) in the collision cell. This method of operation brings advantages to DIA acquisition as, on the one hand, it provides additional ion separation (in the ion mobility dimension), which is important for DIA methods where multiple precursors are fragmented at the same time; and on the other hand, it provides a significant increase in sensitivity by increasing the duty cycle with the addition of the ion mobility dimension to the *m/z* windows schema of the DIA methods.

Many software tools have been created for the processing of DIA runs for quantitative proteomics application, i.e. the quantification of up to thousands of proteins relatively between different groups of samples. Even for diaPASEF, despite its still short lifetime, some tools have been developed or optimized. Among the most widely used tools at present, because they have improved proteome coverage and quantification accuracy, are DIA-NN [5], Spectronaut [6] and the FragPipe suite, which has several tools, including the search engine MSFragger [7], useful for building spectrum libraries. With these tools it is possible to carry out the more traditional workflow, based on a library previously obtained from the identifications in DDA runs, or the more innovative library-free workflow, or more correctly, DDA-based library-free, as in fact an in-silico library is built from a database (DIA-NN) or from the same DIA runs used for quantification (directDIA analysis in Spectronaut).

Nowadays, one of the most popular search engines is MSFragger, which is integrated in the FragPipe proteomics suite. FragPipe, combining MSFragger with the library building tool EasyPQP, makes it possible to build libraries relatively quickly for subsequent use in tools for quantitative analysis of DIA careers. Among these tools, DIA-NN has also gained popularity, currently allowing both a library workflow and a library-free workflow (in which DIA-NN itself, in a stage prior to quantification based on DIA careers, builds a library from a fasta database and the DIA careers). Due to the analysis possibilities offered by FragPipe and DIA-NN, they can be combined in different workflows to perform relative quantitative proteomics: library construction in FragPipe with DDA runs and analysis of DIA runs in DIA-NN [8]; library-free in DIA-NN; with FragPipe, we can even do without DIA runs and conduct an MS1-based label-free quantification (LFQ) with DDA runs using IonQuant [9]. In version 17 of FragPipe, MSFragger version 3.4 was introduced, which offers the possibility to search directly the DIA runs, eliminating the need for DDA runs. This gives us another analysis possibility: building the library with FragPipe from the DIA runs, followed by the quantitative analysis of the same DIA runs in DIA-NN, although at the moment this is not available for diaPASEF runs, only for Sciex (e.g. SWATH) and Thermo DIA files.

Since the first comparative study of software tools for DIA data analysis, specifically Sciex’s SWATH data, was published in 2016 [10], several studies have benchmarked different workflows for DIA quantitative proteomics. Most of them focus on sample preparation strategies and how data are acquired in the mass spectrometer. Thus, Barkovits et al. [2] studied the influence of the type of spectral library used for quantitative analysis of DIA runs in Spectronaut (version 11) software. They included eight libraries of different complexity obtained with two different search engines (Mascot at Proteome Discover, and Pulsar), although they did not apply it to diaPASEF data, but used Thermo Q-Exactive equipment for the acquisition of DDA and DIA runs. Therefore, these results would not be extrapolable to diaPASEF runs obtained in the Bruker timsTOF Pro, where we also have the extra dimension of ion mobility.

In a similar study, also using a Thermo Q-Exactive instrument to obtain DDA and DIA runs, Dowell et al. [11] studied the effect of different types of experimental design and statistical approach on quantitative performance in label-free DDA- and DIA-based quantitative proteomics. Gotti et al. [12] compared combinations of different DIA window acquisition schemes and different software tools (DIA-NN, DIAUmpire, OpenSWATH, ScaffoldDIA, Skyline and Spectronaut) for the analysis of DIA runs acquired on a Thermo Orbitrap instrument. It is therefore noted that the benchmarking studies have focused on DIA data from Thermo’s Orbitrap equipment. It should also be borne in mind that, in general, these benchmarking studies use standard benchmark samples, for example, obtained by mixing commercial standards of protein digests (e.g. HeLa plus *E. coli* digests), or even standard protein mixtures, which therefore suffer from an artificially low biological background variability. Fröhlich et al. [13] did use real clinical samples with higher variance, derived from FFPE lymph node tissue, for comparing four DIA software tools (DIA-NN, Skyline, OpenSwath and Spectronaut) combined with different libraries obtained in-silico or from DDA runs. However, they focused on the analysis of DIA runs obtained, again, with a Thermo Orbitrap mass spectrometer.

Regarding diaPASEF data, Demichev et al. [8] reported the good performance of the combination of building the library in MSFragger and then using this library to analyze the diaPASEF runs in DIA-NN. However, a study comparing the different analysis possibilities offered by the latest software developments optimized for diaPASEF data analysis is still missing.

In this work, we compare the results obtained using different data analysis alternatives that are possible combining the most recent versions of two of the most up-to-date and popular tools, FragPipe and DIA-NN, but applying them to analyze diaPASEF and dda-PASEF runs obtained from a Bruker timsTOF Pro instrument. We compare the results to those obtained with the most up to date version (v15.5) of the gold-standard software for DIA data processing, Spectronaut, and we include library-based and library-free approaches. In addition, since the samples were also run with DDA-PASEF for the library-based workflows, we will include in the comparison the MS1-based LFQ from those runs, using the recently developed IonQuant software with the Match-Between-Runs (MBR) functionality [14], also integrated into FrapPipe. As an illustrative example of application with real samples, such as those that may occur in a real laboratory experiment, we apply the different data analysis workflows to diaPASEF and DDA-PASEF LC-MS runs obtained from the protein extracts of an inflammatory murine cell model, lipopolysaccharide (LPS)-activated macrophages, with and without two anti-inflammatory compounds. The aim is to provide information on how many and which proteins can be quantified, data completeness, precision, and analysis time, so that the end user doing relative quantitative proteomics knows what to expect and what is possible depending on the tool used.

## 2. Methods

### 2.1. Sample collection

RAW 264.7 macrophages (ATCC, TIB-71) were cultured in 24-well plates (Merck) at 37°C and 5% CO2 in sextuplicate, under three different conditions, control (DMEM medium, ATCC-30-2002), anti-inflammatory compound A (DMEM medium with 10 mg/mL of inulin), and antiinflammatory compound B (DMEM medium with 100 μM hydrocortisone). Cells were previously stimulated with Escherichia coli O111:B4 lipopolysaccharides (final concentration 100 ng/ml) during 30 min, and then incubated with treatment for 24 h. Cells from the three groups (LPS, ALPS and B-LPS), six biological replicates each, were collected and lysed, and protein extracts were cleaned by acetone precipitation. Protein pellets were resuspended in 0.2% RapiGest SF (Waters), and protein concentration was measured in a Qubit nanofluorimeter (Thermo Fisher Scientific) using the Qubit Assay Kit (Thermo Fisher Scientific). For each sample, 30 μg of protein were digested with trypsin (Promega) in two steps (2 h at 37 ºC with 1:40 trypsin:protein ratio, plus 15 h after adding another 1:40 trypsin amount) as described elsewhere [15].

### 2.2. LC-MS acquisition

Peptide digests were diluted with 0.1% FA to 100 ng/uL of equivalent protein content. Samples (2 μL, 200 ng protein digest on column) were analyzed on a timsTOF Pro (Bruker) Q-TOF mass spectrometer coupled to a nanoElute (Bruker) LC system. A C18 Aurora Series UHPLC emitter column (250 mm x 75 μm id, 1.6 μm, 120 Å pore size) (IonOpticks) was used for all the analysis, using a trap-elute configuration with an Acclaim PepMap C18 (5 mm, 300 μm id, 5 μm particle diameter, 100 Å pore size) trap cartridge (Thermo Fisher Scientific). The gradient and LC parameters were the same for all analysis: peptides were eluted in a 45 min gradient from 5 to 30% B (from 5 to 25% B in 40 min, from 25 to 30% B in 5 min), plus 12 min of column cleaning (80% B), being A water with 0.1% FA and B ACN with 0.1%FA, respectively. The chromatography flow rate was 300 nL/min. As the peptides eluted from the chromatography to the mass spectrometer, they were ionised in a Captive nano-electrospray source (Bruker) at 1500V.

Samples were run using two different acquisition methods:

(i) for diaPASEF, samples were acquired with a diaPASEF method consisting on 12 cycles including a total of 34 mass width windows (25 Da width, from 350 to 1200 Da) with 2 mobility windows each, making a total of 68 windows (**Supplementary Table S1**) covering the ion mobility range (1/K_0_) from 0.64 to 1.37 V s/cm^2^. These windows were optimized with the Window Editor utility from the instrument control software (timsControl, Bruker) using one DDA-PASEF run acquired from a pool of the analyzed samples. Briefly, this utility loaded the run and represented its ion density in the *m/z* and ion mobility ranges (i.e. the mobility heatmap), so the dia-PASEF windows coverage could be adjusted to ensure complete coverage, and the window settings calculated. The collision energy was programmed as a function of ion mobility, following a straight line from 20 eV for 1/K_0_ of 0.6 V s/cm^2^ to 59 eV for 1/K_0_ of 1.6 V s/cm^2^. The TIMS elution voltage was linearly calibrated to obtain 1/K_0_ ratios using three ions from the ESI-L Tuning Mix (Agilent) (*m/z* 622, 922, 1222) before each run, using the ‘Automatic calibration’ utility in the control software (timsControl, Bruker). One of the samples (A-LPS-1) was run in quintuplicate.

(ii) additionally, samples were also acquired with a DDA-PASEF method consisting on 10 MS/MS PASEF scans per topN acquisition cycle, with an accumulation time of 100 ms and a ramp of 100 ms. MS and MS/MS spectra were acquired in an *m/z* range from 100 to 1700 and in an ion mobility range (1/K_0_) from 0.60 to 1.60 V s/cm^2^, selecting precursor ions for the MS/MS PASEF scans from a previous TIMS-MS scan. The collision energy was programmed as a function of ion mobility, following a straight line from 20 eV for 1/K_0_ of 0.6 to 59 eV for 1/K_0_ of 1.6. The TIMS elution voltage was linearly calibrated to obtain 1/K_0_ ratios using three ions from the ESI-L Tuning Mix (Agilent) (*m/z* 622, 922, 1222). One of the samples (A-LPS-1) was run in quintuplicate. Equal aliquots from the six biological replicates for each group were pooled in a vial, and the three pools were also analyzed with this DDA-PASEF method.

### 2.3 Data analysis workflows

A total of eight different workflows were run (**Figure 1**), as described hereafter. All analysis were run on an AMD Ryzen Threadripper 3970X 32-core processor (64 logial CPU cores), with 128 Gb RAM, running in Windows 10 (AMD64 architecture) as OS and Java version 1.8.0. All files used and generated, including intermediate temporary files, were loaded from and saved to a 7200 rpm hard drive disk.

**Figure 1.**
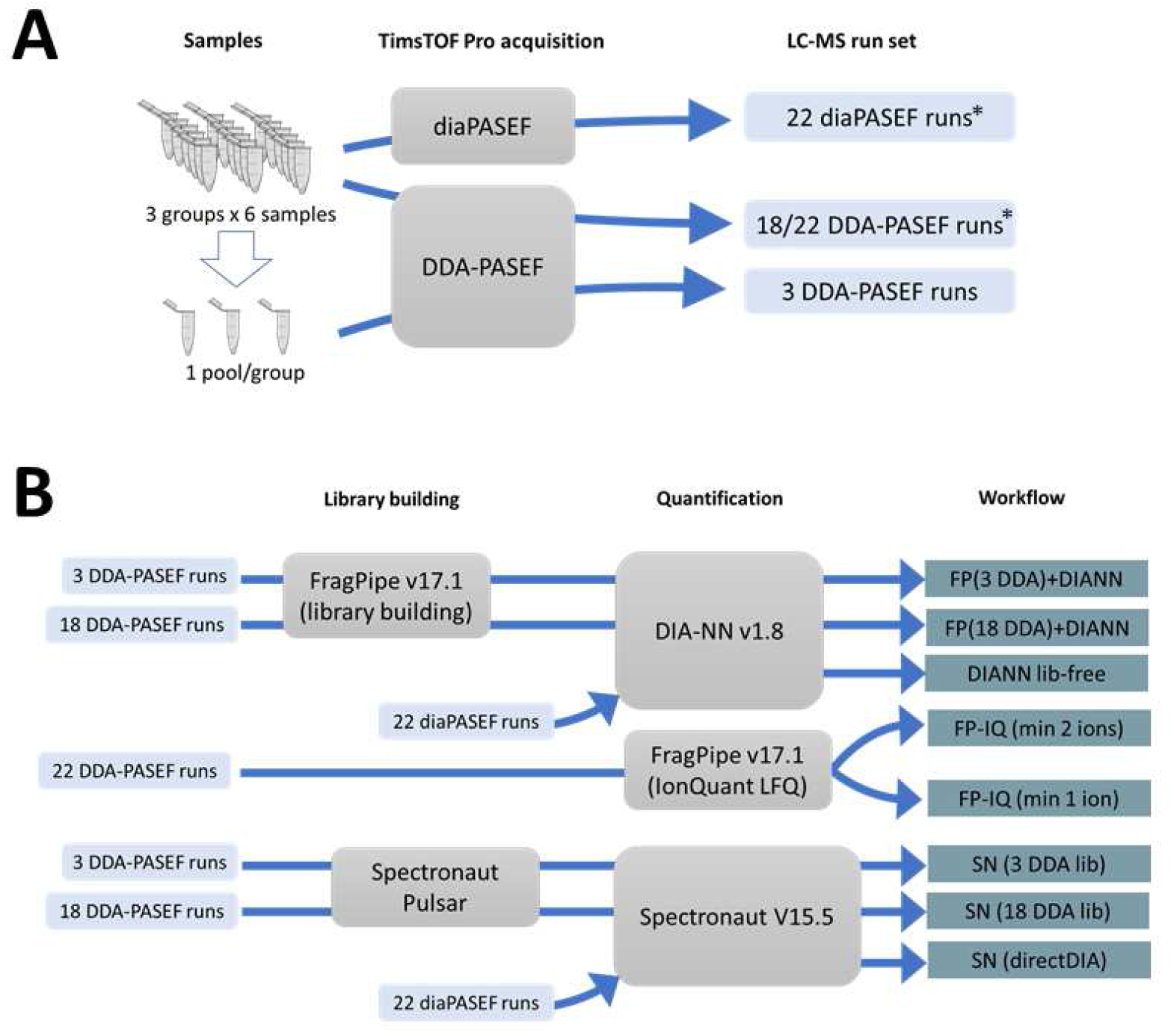
Study design for (**A**) sample acquisition using either diaPASEF or DDA-PASEF, and (**B**) data analysis for the different workflows. (*) The diaPASEF and DDA-PASEF run set used for quantification included 4 injection replicates of one of the samples, thus 22 runs instead of the 18 runs used for library building.

#### 2.3.1. Library generation in FragPipe and diaPASEF analysis in DIA-NN

Two different libraries were tested, one built from the 18 DDA-PASEF runs, and the other built with 3 DDA-PASEF runs from the 3 pooled vials obtained after mixing an aliquot of the samples from each of the 3 groups. Both libraries were built in the same way: the workflow ‘DIA-SpecLib_Quant’ from FragPipe version 17.1 was used, including database search with MSFragger (version 3.4), deep-learning prediction rescoring with MSBooster, Percolator and ProteinProphet (Philosopher version 4.1.0) for PSM validation and protein inference, and EasyPQP (version 0.1.25) for building the spectral library, unchecking the option ‘Quantify with DIA-NN’. The raw .d files from Bruker were searched against a mouse (UP000000589) isoform fasta database (downloaded on 09/07/2021) appended with common contaminant proteins, containing in total 25483 sequence entries. Decoy reversed sequences were appended to the search database. The default MSFragger search parameters were used, except precursor and fragment mass tolerances were set to 20 ppm; enzyme: trypsin (not cutting before P); peptide length 7-30. Variable modifications were set to oxidation of Met and acetylation of protein N-t, and fixed modifications to carbamidomethylation of Cys. MSBooster, Percolator and ProteinProphet default options were used, and results were filtered by 1% FDR at protein level. The spectral library was generated with EasyPQP, with the automatic selection of a run as reference for RT and ion mobility calibration. The two libraries thus produced were used in DIA-NN version 1.8 to analyze the diaPASEF runs. In DIA-NN, missed cleavages were set to 0, precursor change range 2-4, and precursor *m/z* range 349-1500, neural network classifier set to double-pass mode, quantification strategy was set to ‘Robust LC (high precision)’, and MBR option was enabled. MS1 and MS2 accuracy, and retention time window scans, were set to 0 in order to let DIA-NN to perform their automatic inference for the first run in the experiment. All other DIA-NN settings were left default, using RT-dependent cross-run normalization and filtering the output at 1% FDR. The number of threads used by DIA-NN, were 32, as automatically suggested by the software. In order to quantify the effect of the MSBooster application, recently implemented into FragPipe v17.1, these two analyses were repeated in exactly the same way, but obtaining the spectral libraries with the immediately previous version of FragPipe (v16.0), in which the MSBooster application did not yet exist.

#### 2.3.2. DDA-library free diaPASEF analysis in DIA-NN

DIA-NN version 1.8 was used first to build an in-silico predicted library from the same fasta database (mouse UP000000589), enabling the options ‘FASTA digest for library-free search/library generation’ and ‘Deep learning-based spectra, RTs and IMs prediction’, with the rest of parameters in the Precursor ion generation section as above; and, on a second step, this predicted library was used to analyze the diaPASEF dataset, with the same parameters as above.

#### 2.3.3. Spectronaut, library-based

Two libraries were generated using Spectronaut (v15.5) with Pulsar search engine, one built from the 18 DDA-PASEF runs, and the other built from the 3 DDA-PASEF runs from the pooled vials obtained from mixing an aliquot of the samples from each of the 3 groups. The same database (mouse UP000000589 isoform fasta database, downloaded on 09/7/2021, containing 25367 sequences) as before was used for the search, together with a fasta file containing 112 common contaminant sequences. The default factory settings were used for the Pulsar search and library generation (including Trypsyin/P as enzyme, 7-52 peptide length range, up to two missed cleavages allowed, Oxidation of Me and acetylation of Protein N-t as variable modifications, carbamidomethyl of C as fixed modification, and 1% FDR for PSM, peptide and protein identification). The two libraries generated were used to carry out two DIA analysis with Spectronaut (v15.5). The default factory settings were used, except for the calibration MS1 and MS2 mass tolerances, which were set to 20 ppm; proteotypicity filter, set to ‘only protein group specific’; and no imputing strategy. Local cross run normalization strategy and MaxLFQ method were used for protein quantification, and quantity was determined at MS2 level using the area of extracted chromatogram traces.

#### 2.3.4. Spectronaut, library-free (directDIA)

The directDIA workflow in Spectronaut (v15.5) was used for analyzing the diaPASEF dataset with no need to build a library from the DDA runs. The settings for Pulsar search and DIA analysis steps were the same as in the Spectronaut library-based workflow above.

For comparison, we repeat the same analyses, with the same settings, on the immediately preceding Spectronaut version (v15.4).

#### 2.3.5. MS1 LFQ quantification with Fragpipe IonQuant

The DDA-PASEF individual runs were analyzed in Fragpipe version 17.1 with the ‘LFQ-MBR’ workflow, which included database search with MSFragger (version 3.4), deep-learning prediction rescoring with MSBooster, Percolator and ProteinProphet (Philosopher version 4.1.0) for PSM validation and protein inference, and MS1-level label-free quantification with IonQuant. All parameters for MSFragger, MSBooster and Philosopher were the same as those used in the ‘library generation in Fragpipe’ workflow above. For MS1-leve quantification in IOnQuant, MBR was enabled, and the MaxLFQ algorithm was allowed, running two different analyses, with 2 or 1 minimum number of ions required for quantifying a protein. For feature detection in IonQuant, 10 ppm, 0.4 min, and 0.05 1/K_0_ were set for *m/z*, RT and IM tolerances, respectively.

## 3. Results and discussion

Three run sets were obtained from the 18 samples, comprising: 22 diaPASEF runs (one run per sample, plus four more injection replicates for sample A-LPS-1); 22 DDA-PASEF runs (same as for the diaPASEF runs); and 3 DDA-PASEF runs (one run per group pooled vial).

Quantitative reports were obtained from the eight data processing workflows, and different metrics of interest to the end-user were calculated. The aim is not to compare quantification accuracy using standard digest mixtures such the two-organism benchmark datasets, but to provide the potential user with information on what can be expected and what is possible to obtain depending on the tool used, in terms of how many and which proteins can be quantified, data completeness, precision, and computing time, using a real-life sample set like the one that can be found in any proteomics experiment.

First, we compared the reported quantitative values for the protein groups quantified in one of the samples (A-LPS-1) by pairs of workflows, calculating Pearson’s correlation coefficients. It can be seen from **Supplementary Figure S1** that the eight workflows correlated well with each other, with all Pearson correlation coefficients above 0.69. As expected, correlation was higher within related workflows (0.87 to 0.92 between DIA-NN based workflows; 0.89 to 0.91 between Spectronaut analyses), but was also higher between DIA-NN and Spectronaut workflows (0.85 to 0.89). The lowest correlation was found between IonQuant quantifications and Spectronaut analyses (0.69 to 0.62).

**Figure 2** shows useful indicators for comparing the different data-analysis workflows, e.g. quantified protein groups, data completeness and precision.

**Figure 2.**
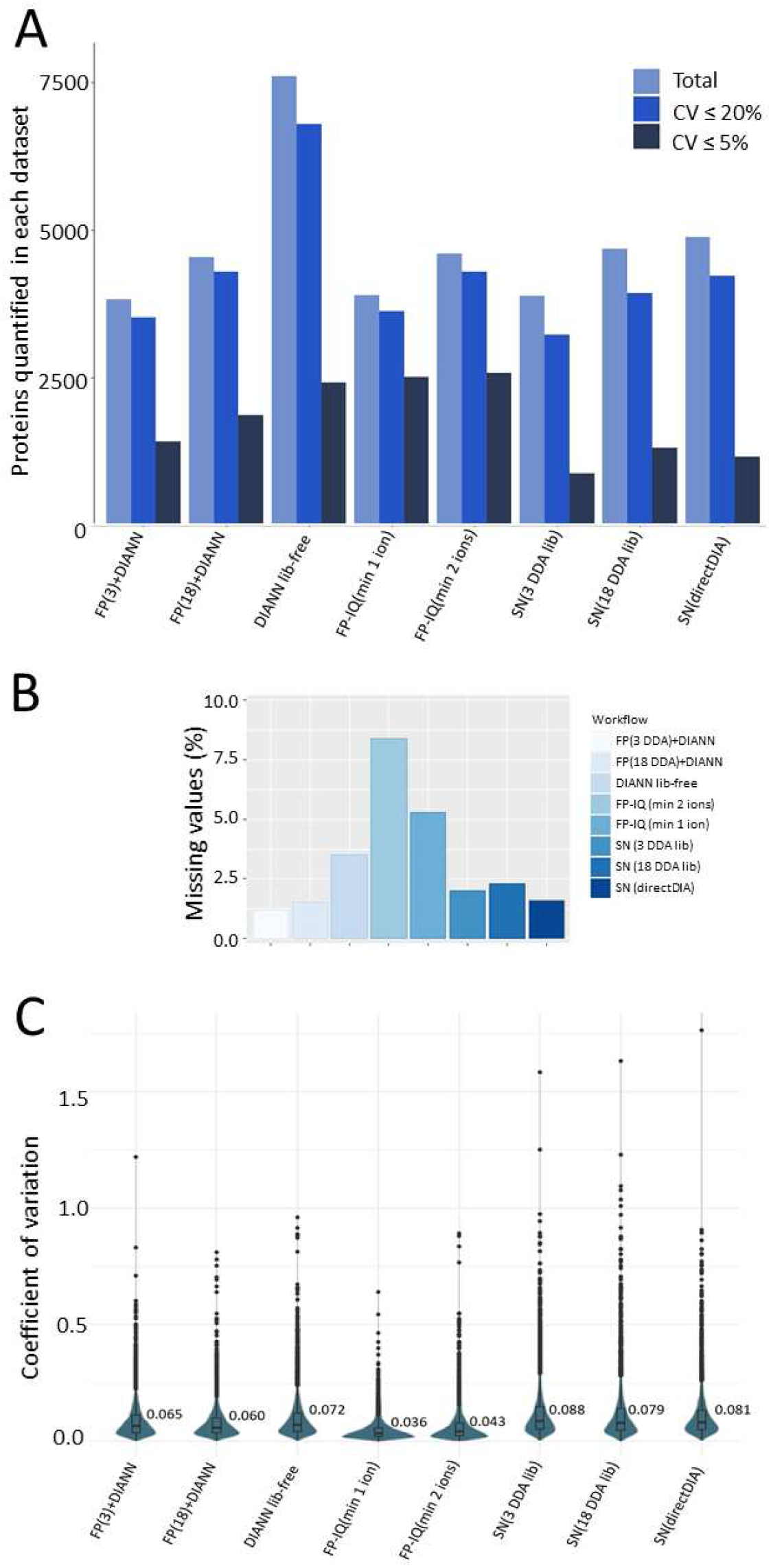
Quantified protein groups, data completeness and precision of the eight data-analysis workflows. (**A**) Total number of protein groups quantified in each workflow, showing also the number of those with coefficient of variation (CV) up to 20% and up to 5%. (**B**) Percentage of missing values obtained for each dataset. (**C**) CVs of the quantified proteins, showing the median CV in the inserts.

Protein groups quantified in each individual dataset ranged from 3823 when using DIA-NN with the smallest FragPipe built library, to 7606 protein groups in library-free DIA-NN (**Figure 2A**), with the number of proteins quantified with 20% and 5% CV thresholds varying accordingly. Notably, a majority of the quantified proteins showed a CV below 20%: over 90% in the DIA-NN or IonQuant workflows, and between 83 and 87% in the Spectronaut workflows.

Far from the 7606 protein groups quantified by DIA-NN only, in the mid-range is the library-free workflow of Spectronaut (directDIA, with 4875 protein groups), very similar to the workflows based on the large library (4679 in Spectronaut, 4543 in FragPipe+DIA-NN) and the IonQuant LFQ quantification using one minimum ion (4595 protein groups). Trailing behind are IonQuant LFQ with 2 minimum ions (3897 protein groups) and the analyses using the smallest libraries (3877 in Spectronaut, 3823 in FragPipe+DIA-NN). All runs where single-shot, that is, no fractionation was made for either the diaPASEF runs or the DDA-PASEF runs. This is important to note as library-based workflows would benefit, in the number of proteins in the library, and therefore in the number of proteins quantified, from obtaining the library after sample fractionation. However, this would be at the cost of increasing equipment usage time and cost by increasing the number of DDA runs.

Within the workflows in which we used FragPipe, one of the processing steps is the validation of the MSFragger search results with Philosopher, and at this stage the MSBooster for deep learning-based rescoring has been introduced in the latest version of FragPipe (v17.1). In order to estimate the effect of MSBooster on the number of proteins quantified, we repeated both FP+DIA-NN analyses using FragPipe v16.0 (without MSBooster), quantifying 3451 and 4060 protein groups (for FP(3 DDA)+DIANN and FP(18 DDA)+DIANN workflows), instead of the 3823 and 4543 that were quantified with version 17.1, already including MSBooster. This means that FragPipe, in its new version, has made a quantitative leap, which we quantify as 10% more protein groups quantified.

In addition to the number of proteins quantified in the dataset, the completeness of each dataset is an important indicator. The percentage of missing data with respect to the total number of measurements in each dataset is shown in **Figure 2B**. The highest data completeness (98.8% and 98.5%) was achieved by the FragPipe+DIA-NN workflows. These results improve on those reported by Demichev et al. [8], where data completeness ranged from 94% to 98%. On the other hand, LFQ with IonQuant using 2 min ions, resulted in the highest percentage of missing values (8.4%). The percentage decreased to 5.4% when using 1 min ion in IonQuant, although it was still higher than that obtained by the other tools. Spectronaut achieved good data completeness, with 2 and 2.3% missing data for the library-based workflows, and even less (1.6%) for the directDIA analysis. Library-free DIA-NN results are in the mid-range, with a 3.5% of missing values, which is not an excessive cost in exchange for the large number of proteins it quantifies.

As an indicator of each workflow quantitative precision, the coefficients of variation (CV) of the quantitative values of each quantified protein group were calculated using the five injection replicates of the sample A-LPS-1, and are shown for the eight workflows in **Figure 2C**. Median CVs were below 9% in all workflows, with Spectronaut in the range 7.9 to 8.8%, and label free DIA-NN with 7.2%. Library-based DIA-NN median CVs obtained (6.5% and 6%) are in the range of those reported by Demichev et al. [8] (2.7% to 9.5%). FragPipe’s IonQuant performed the best in terms of median CV (3.6% and 4.3% for the 2 and 1 minimum ion analysis, respectively). Notably, these CVs obtained by IonQuant were better than those reported by Yu et al. [9] in the original IonQuant publication (6.4 and 7.3% median CVs), where FDR-controlled MBR was not available, but consistent with what was reported by the same authors following MBR implementation (3.6 and 4.0% median CVs) [14].

Within the two IonQuant options tested, the choice of 1 minimum ion resulted in more quantified protein groups than taking 2 minimum ions (4595 vs 3897), but at the cost of the CVs going up, although it is a slight increase and the difference in quantified proteins is so large that the number of quantified protein groups with CV threshold of 20% is still higher (4288 for 1 minimum ion vs. 3617 for 2 minimum ions). Therefore, even with a higher median CV, the 1 minimum ion option would be recommended over the 2 minimum ion one.

In the Spectronaut workflows, the median CVs were higher, and there was also a lower proportion of quantified proteins with CV less than 5% (**Figure 2A**). Within the Spectronaut workflows, it was directDIA that quantifies the most protein groups with CV<20%, in the range of what library-based FragPipe+DIA-NN and LFQ FragPipe-IonQuant analyses achieved, although if we go down to a threshold of 5% CV, directDIA is slightly surpassed by FragPipe-IonQuant (despite quantifying more than 3000 protein groups less).

For simplicity of analysis, the possibility of semi-tryptic peptides due to gas-phase fragmentation, prior to MS/MS fragmentation, which has been reported to occur frequently in timsTOF equipment [9], was not taken into account in the parameters used. Nevertheless, although an increase in the number of proteins identified when performing a semi-enzymatic search has been reported, it was found that it did not increase the number of proteins quantified, and that taking this into account in the data analysis does not seem to be beneficial [8].

Next, we checked how many proteins were quantified together in various workflows, and the number of proteins that were quantified exclusively in a specific workflow (**Figure 3**). When comparing the analyses performed with DIA-NN (library free and FragPipe-built library with either 3 or 18 DDA runs) (**Figure 3A**), a central core of 2958 protein groups quantified by the 3 analyses was found, but the DIA-NN library-free analysis alone contributed 3876 more. To note, even the DIA-NN analysis with the smallest library (FP(3 DDA)+DIANN) accounted for a set of 170 protein groups that was not quantified by the other two DIA-NN workflows. There were 260 protein groups (170 + 90) quantified by this 3-DDA library-based DIA-NN that were not quantified by DIA-NN using the largest library. And vice versa, 980 (682+298) protein groups quantified by DIA-NN with the large library, which are not quantified with the library built from the 3 DDA-PASEF run set. However, in the LFQ quantification with IonQuant, the workflow with the most protein groups quantified (the one using 1 min ion) quantifies all those quantified by the 2 min ion workflow (3897), plus 698 more (**Figure 3B**). For the Spectronaut options (**Figure 3C**), the largest group is that of the proteins quantified in the three workflows (3480), the nucleus on which directDIA analysis contributed 1395 (689+623+83) protein groups more, 623 of them exclusively. When comparing all workflows together (**Figure 3D**), it is observed that a central core of 2643 protein groups were quantified in all analyses, but those quantified exclusively by library-free DIA-NN are even more (2669).

**Figure 3.**
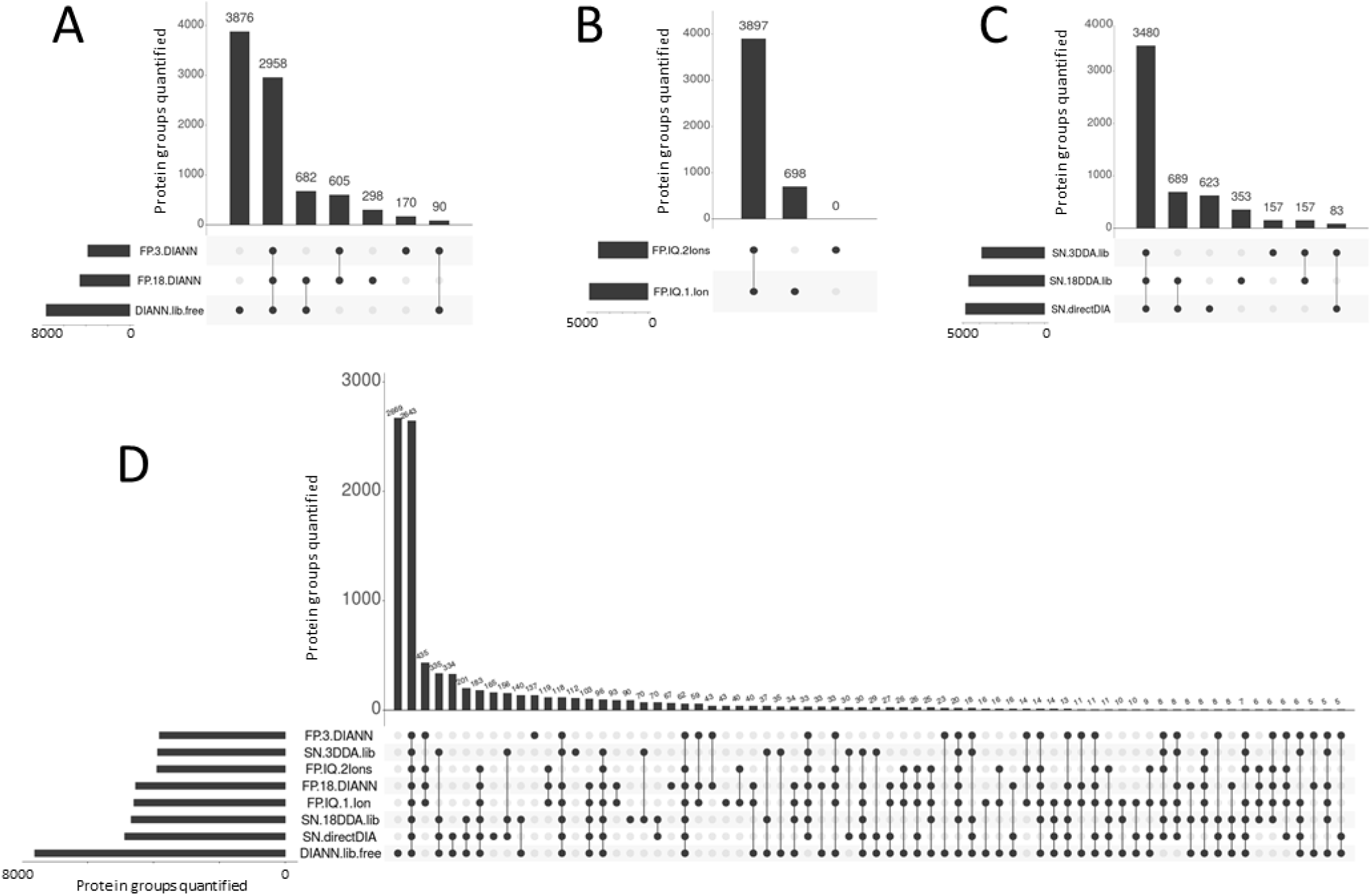
Comparison of protein groups quantified by each data-analysis workflow, showing (**A**) DIA-NN workflows only; (**B**) IonQuant workflows only; (**C**) Spectronaut workflows; and (**D**) all workflows together.

Finally, we compared the computational time spent for each analysis. In **Figure 4** these running times, together with the number of protein groups quantified, are shown for each data analysis workflow.

**Figure 4.**
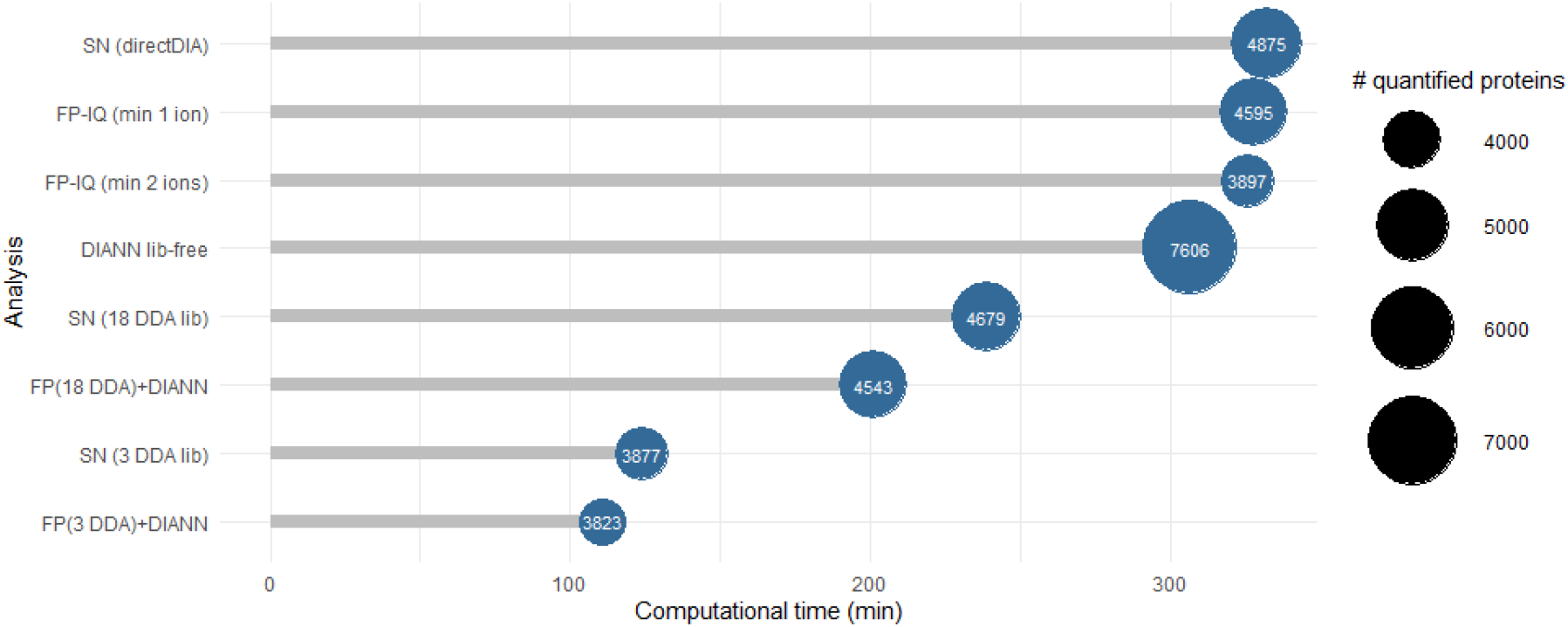
Computing time spent on the different analysis, indicating the number of protein groups quantified in each dataset.

As expected, in workflows with spectral library construction, the time taken by both FragPipe and Spectronaut depends on the library size, and thus the number of DDA runs used to perform the search, with the total time doubling from three to 18 DDA runs used, although, as expected, this is accompanied by an increase in the number of proteins quantified, which is remarkable: the number of protein groups quantified is increased by 19% for FragPipe + DIANN, and by 21% for Spectronaut. In any case, time taken by library-free workflows is even longer (slightly above 5 hours), despite the fact that fewer runs are processed. On the other hand, these library-free flows were able to quantify more proteins than library-based ones, especially the DIA-NN analysis with 7606 protein groups quantified.

As Spectronaut developers claimed after releasing the software version 15.5, they have significantly improved the processing speed for the directDIA analysis compared to previous versions: when we repeated the analysis in the immediately preceding (v15.4), computing speed was significantly faster in the new version (431 minutes in v15.4 to 332 minutes in v15.5, i.e. 23% less time); the number of protein groups quantified, however, increased in the new version by 3.2% (from 4722 to 4875). This improvement makes Spectronaut directDIA workflow very competitive in terms of speed and quantified proteins.

The duration of IonQuant LFQ quantification in FragPipe is in the upper range (above 5 hours), without providing more quantified proteins than faster workflows based on the largest libraries. It is important to note that in addition to the data analysis computation time reported here, we have not considered the equipment usage time, and economic cost if any, of using one or the other workflows. In our example application, we would move from 18 diaPASEF runs (in the case of directDIA or DIA-NN) or DDA-PASEF runs (LFQ with FragPipe-IonQuant) to 36 runs (18 diaPASEF runs plus the 18 DDA-PASEF runs for the largest library) for the DDA-library analyses, thus doubling the equipment time and economic cost of the analysis. Consequently, in relation to these indicators, equipment time and cost, the library-free designs, acquiring only one diaPASEF (directDIA, library-free DIA_NN) or DDA-PASEF (LFQ with FragPipe IonQuant) run per sample are more efficient than the library-based workflows. We have verified here that library-independent workflows, not so efficient not so long ago, have greatly improved in the recent versions of the software tools, and now perform as well or even better than library-based ones. Therefore, only in case equipment usage time and cost are not limiting we can recommend library-based workflows.

For a fast visual comparison of the results of the eight workflows, we displayed each variable to compare (number of protein groups quantified, number of protein groups quantified with CV ≤ 20% in the five replicates, median CV, data completeness, computing time, and number of LC-MS runs needed (and thus cost)) in a radar plot (**Figure 5**). All values within a variable were rescaled in a score range from 0 to 1, with 1 for the maximum of the variable in the eight workflows (e.g. 7606 protein groups in the DIA-NN workflow), and 0 for the minimum (e.g. 3823 protein groups in FP(3 DDA)+DIANN). For variables where less is better (e.g. computation time, median CV, runs & cost), rescaling was applied to the inverse of the raw value. In this way, the better the performance of a variable (e.g. 7606 protein groups in the DIA-NN analysis; faster computation time), the further out of the radar plot that variable is represented; and vice versa, when the result is worse, the further inside the radar plot the point corresponding to that variable is represented (e.g. 3823 protein groups in FP(3 DDA)+DIANN; longer computation duration). In addition to the profile of every workflow (blue line), an average profile of the eight workflows (grey shading) is also shown in each individual plot. For instance, we can see that DIA-NN is the workflow with the best result in terms of total quantified proteins, far above average; or that IonQuant with 2 minimum ions has the best result in median CV; or quickly check which workflows perform best in data completeness.

**Figure 5.**
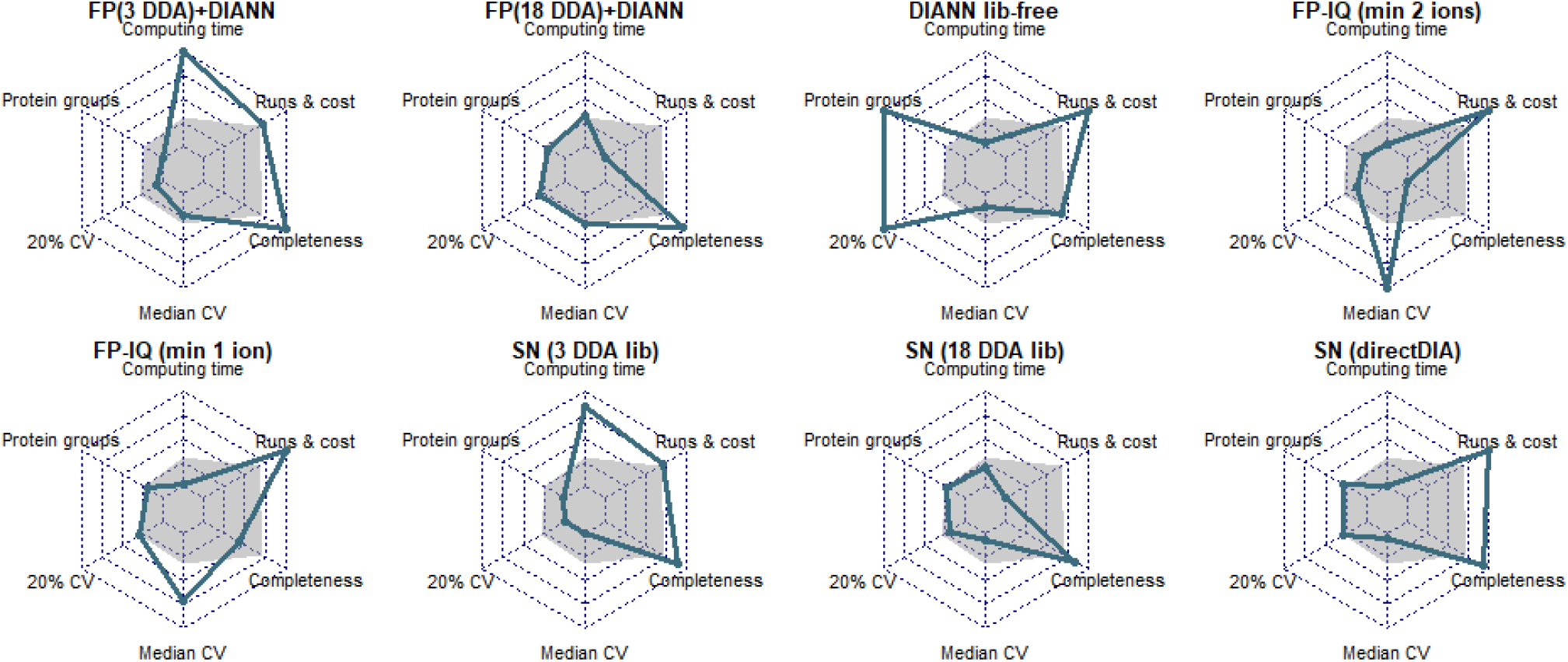
Radar plots for the eight workflows showing the performance on six variables (blue dots): number of protein groups quantified, number of protein groups quantified with CV ≤ 20% in the five replicates, median CV, data completeness, computing time, and number of LC-MS runs needed (and cost). The values of each variable were rescaled between 0 (worst value of the variable, more towards the center of the plot) and 1 (best result, the more outside on the plot). An average profile of the eight workflows is shown with grey shading on each plot.

## 4. Conclusion

In this paper we compare the results of analyzing PASEF data with different combinations of library-based and library-free analysis, combining the tools of the FragPipe suite, DIA-NN and including MS1-level LFQ with DDA-PASEF data, and comparing with the workflows possible in Spectronaut. We found that library-free workflows have experienced an increase in efficiency, both in number of proteins identified and in computational speed, with similar or even better performance than library-based and MS1-level LFQ workflows in terms of number of proteins quantified, precision (coefficient of variation) and data completeness. Therefore, only in case equipment usage time and cost are not limiting we can recommend library-based workflows, and even in that case, we have found that library-free workflows such as the one performed by DIA-NN greatly increase the number of proteins quantified without excessive time costs. On the other hand, Fragpipe’s LFQ of DDA-PASEF runs with IonQuant, although it manages to quantify a lower number of proteins than the library-free flow in DIA-NN, has the best values in coefficients of variation, being at the level of library-based DIA analyses in terms of quantified proteins.

In conclusion, we report here information so that the user who is going to conduct a relative quantitative proteomics study using a timsTOF Pro mass spectrometer can make an informed decision on how to acquire (diaPASEF for DIA analysis, or DDA-PASEF for MS1-level LFQ) his samples, and which data analysis software and workflow to use, among the different alternatives offered by the recently optimized tools for TIMS-PASEF data analysis.

## SUPPLEMENTARY MATERIAL

**Supplementary Table S1.** Windows scheme used for the diaPASEF acquisition. IM, ion mobility.

| MS Level | TIMS Scan | Start Mass [m/z] | End Mass [m/z] | Start IM [1/K0] | End IM [1/K0] |
| --- | --- | --- | --- | --- | --- |
| MS1 | 0 | na | na | na | na |
| dia-PASEF | 1 | 1125 | 1150 | 1.25 | 1.34 |
| dia-PASEF | 1 | 975 | 1000 | 1.15 | 1.24 |
| dia-PASEF | 1 | 825 | 850 | 1.05 | 1.14 |
| dia-PASEF | 1 | 675 | 700 | 0.95 | 1.04 |
| dia-PASEF | 1 | 525 | 550 | 0.85 | 0.94 |
| dia-PASEF | 1 | 375 | 400 | 0.75 | 0.84 |
| dia-PASEF | 1 | 350 | 375 | 0.64 | 0.73 |
| dia-PASEF | 2 | 1100 | 1125 | 1.23 | 1.32 |
| dia-PASEF | 2 | 950 | 975 | 1.13 | 1.22 |
| dia-PASEF | 2 | 800 | 825 | 1.03 | 1.12 |
| dia-PASEF | 2 | 650 | 675 | 0.93 | 1.02 |
| dia-PASEF | 2 | 500 | 525 | 0.83 | 0.92 |
| dia-PASEF | 2 | 350 | 375 | 0.73 | 0.82 |
| dia-PASEF | 3 | 1150 | 1175 | 1.26 | 1.36 |
| dia-PASEF | 3 | 1000 | 1025 | 1.16 | 1.26 |
| dia-PASEF | 3 | 850 | 875 | 1.06 | 1.16 |
| dia-PASEF | 3 | 700 | 725 | 0.96 | 1.06 |
| dia-PASEF | 3 | 550 | 575 | 0.86 | 0.96 |
| dia-PASEF | 3 | 400 | 425 | 0.76 | 0.86 |
| dia-PASEF | 3 | 375 | 400 | 0.65 | 0.75 |
| dia-PASEF | 4 | 1175 | 1200 | 1.28 | 1.37 |
| dia-PASEF | 4 | 1025 | 1050 | 1.18 | 1.27 |
| dia-PASEF | 4 | 875 | 900 | 1.08 | 1.17 |
| dia-PASEF | 4 | 725 | 750 | 0.98 | 1.07 |
| dia-PASEF | 4 | 575 | 600 | 0.88 | 0.97 |
| dia-PASEF | 4 | 425 | 450 | 0.78 | 0.87 |
| dia-PASEF | 4 | 400 | 425 | 0.67 | 0.76 |
| dia-PASEF | 5 | 1050 | 1075 | 1.2 | 1.29 |
| dia-PASEF | 5 | 900 | 925 | 1.1 | 1.19 |
| dia-PASEF | 5 | 750 | 775 | 1 | 1.09 |
| dia-PASEF | 5 | 600 | 625 | 0.9 | 0.99 |
| dia-PASEF | 5 | 450 | 475 | 0.8 | 0.89 |
| dia-PASEF | 5 | 425 | 450 | 0.69 | 0.78 |
| dia-PASEF | 6 | 1075 | 1100 | 1.21 | 1.31 |
| dia-PASEF | 6 | 925 | 950 | 1.11 | 1.21 |
| dia-PASEF | 6 | 775 | 800 | 1.01 | 1.11 |
| dia-PASEF | 6 | 625 | 650 | 0.91 | 1.01 |
| dia-PASEF | 6 | 475 | 500 | 0.81 | 0.91 |
| dia-PASEF | 6 | 450 | 475 | 0.7 | 0.8 |
| dia-PASEF | 7 | 1075 | 1100 | 1.12 | 1.21 |
| dia-PASEF | 7 | 925 | 950 | 1.02 | 1.11 |
| dia-PASEF | 7 | 775 | 800 | 0.92 | 1.01 |
| dia-PASEF | 7 | 625 | 650 | 0.82 | 0.91 |
| dia-PASEF | 7 | 475 | 500 | 0.72 | 0.81 |
| dia-PASEF | 8 | 1100 | 1125 | 1.13 | 1.23 |
| dia-PASEF | 8 | 950 | 975 | 1.03 | 1.13 |
| dia-PASEF | 8 | 800 | 825 | 0.93 | 1.03 |
| dia-PASEF | 8 | 650 | 675 | 0.83 | 0.93 |
| dia-PASEF | 8 | 500 | 525 | 0.74 | 0.83 |
| dia-PASEF | 9 | 1125 | 1150 | 1.15 | 1.25 |
| dia-PASEF | 9 | 975 | 1000 | 1.05 | 1.15 |
| dia-PASEF | 9 | 825 | 850 | 0.95 | 1.05 |
| dia-PASEF | 9 | 675 | 700 | 0.85 | 0.95 |
| dia-PASEF | 9 | 525 | 550 | 0.75 | 0.85 |
| dia-PASEF | 10 | 1150 | 1175 | 1.17 | 1.26 |
| dia-PASEF | 10 | 1000 | 1025 | 1.07 | 1.16 |
| dia-PASEF | 10 | 850 | 875 | 0.97 | 1.06 |
| dia-PASEF | 10 | 700 | 725 | 0.87 | 0.96 |
| dia-PASEF | 10 | 550 | 575 | 0.77 | 0.86 |
| dia-PASEF | 11 | 1175 | 1200 | 1.18 | 1.28 |
| dia-PASEF | 11 | 1025 | 1050 | 1.08 | 1.18 |
| dia-PASEF | 11 | 875 | 900 | 0.98 | 1.08 |
| dia-PASEF | 11 | 725 | 750 | 0.88 | 0.98 |
| dia-PASEF | 11 | 575 | 600 | 0.79 | 0.88 |
| dia-PASEF | 12 | 1050 | 1075 | 1.1 | 1.2 |
| dia-PASEF | 12 | 900 | 925 | 1 | 1.1 |
| dia-PASEF | 12 | 750 | 775 | 0.9 | 1 |
| dia-PASEF | 12 | 600 | 625 | 0.8 | 0.9 |

**Supplementary Figure S1.**
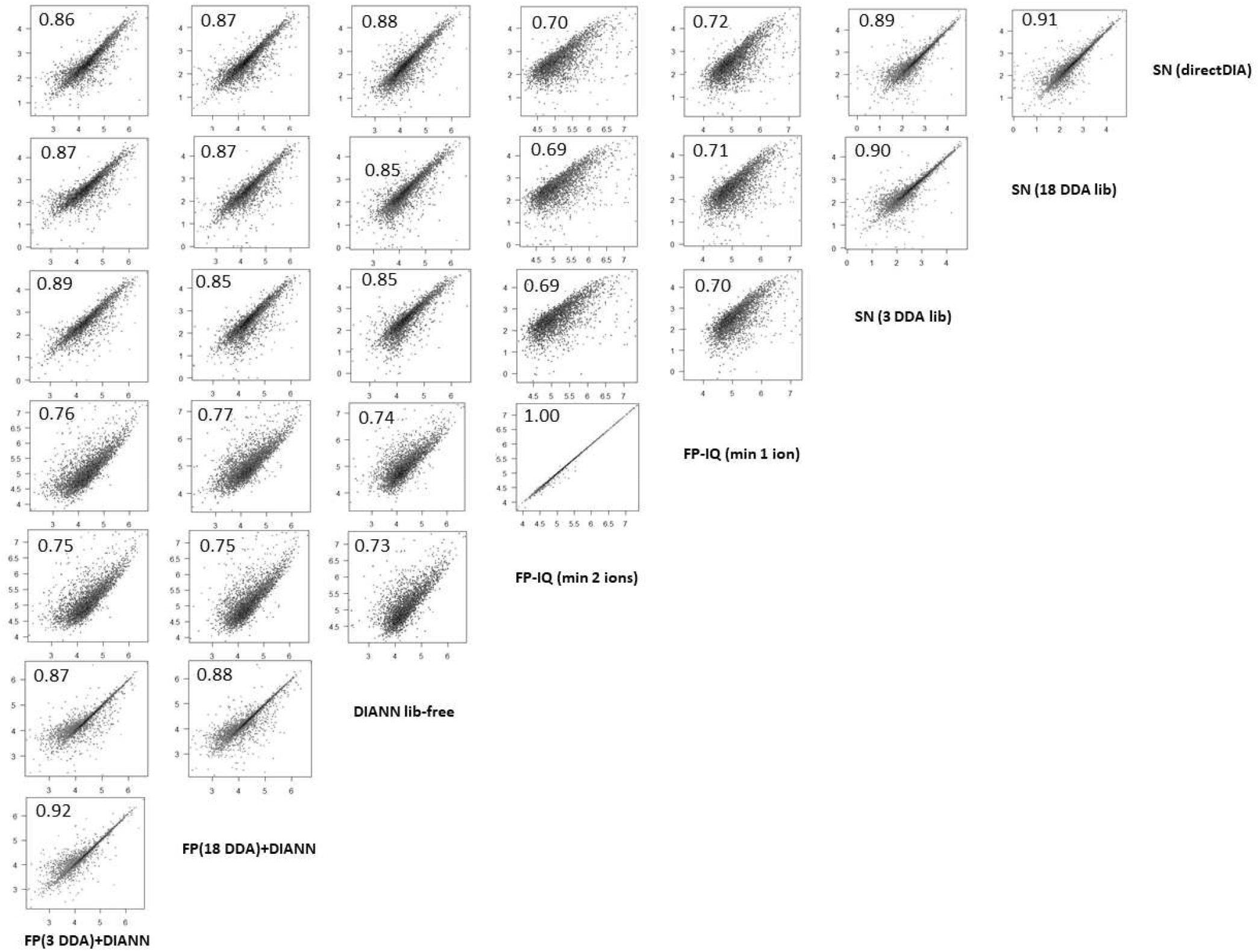
Correlation of quantified protein groups between the different data analysis workflows. Log-transformed abundances from sample A-LPS-1 were used. The Pearson correlation coefficients are shown in the top left-hand corner of each plot.

